# Modifications of the 22A apoA-I mimetic peptide sequence improve the anti-atherosclerotic properties of synthetic HDL

**DOI:** 10.1101/2025.08.13.670037

**Authors:** Sirine Nouri, Laura Giorgi, Akseli Niemelä, Juho Heininen, Khalfa Benadouda, Sagar Dhakal, Artturi Koivuniemi

## Abstract

Synthetic high-density lipoprotein (sHDL) constituted of apolipoprotein A-I (apoA-I) mimetic peptides and phospholipids are nanometer-scale particles designed to recreate biological functions of HDL particles in the context of cardiovascular disease. Particularly, the therapeutic efficacy of sHDL particles is attributed to their ability to promote reverse cholesterol transport (RCT), a process where accumulated cholesterol is transported from peripheral tissues to the liver for elimination. Here, we designed two novel apoA-I mimetic peptides (22A-F and 22A-P-18A) by modifying the sequence of the well-studied apoA-I mimetic peptide 22A. These modifications were intended to improve cholesterol efflux from macrophages *in vitro* and structural stability of sHDL particles in human plasma while preserving their ability to activate lecithin-cholesterol acyltransferase (LCAT). We performed a systematic examination of the potency of sHDL particles made with these peptides in cellular cholesterol efflux, activation of LCAT, plasma HDL remodeling and proteolytic stability. Our study highlights that these modifications improve cholesterol efflux and, in the case of 22A-P-18A, also cholesterol esterification rate by LCAT but they do not appear to influence HDL remodeling in human plasma. Nonetheless, the LCAT activity assay conducted in human plasma suggest that intact sHDL particles are present and are likely the primary contributors to the increased cholesterol esterification rate, rather than the pre-β HDL fraction generated through HDL remodeling. These findings offer new mechanistic insights into how specific peptide modifications affect key steps in RCT, laying the groundwork for future studies to explore their functional relevance in atherosclerosis and HDL-based drug delivery applications.

## Introduction

High density lipoproteins (HDL) are macromolecular complexes of various lipids and apolipoproteins that are known to have important anti-atherosclerotic functions, notably through the enhancement of reverse cholesterol transport (RCT), the process by which excess cellular cholesterol is removed from peripheral tissues and delivered to the liver for excretion in the bile (1–3). Cholesterol efflux is the first step of RCT, where lipid-poor apolipoprotein A-I (apoA-I) particles acquire phospholipids and unesterified cholesterol from cells via the ABCA1 transporter. This step is critical for maintaining intracellular cholesterol homeostasis and for preventing the progression of atherosclerotic plaque, where efficient cholesterol removal from foam cells in the arterial wall is essential (4,5). Cholesterol efflux leads to the formation of discoidal pre-β HDL particles that interact with lecithin-cholesterol acyltransferase (LCAT), an enzyme that facilitates the esterification of cholesterol within HDL. The formed cholesteryl esters, that are more hydrophobic than cholesterol, migrate to the core of pre-β HDL particles, promoting their maturation into spherical mature α-HDL particles (6). Cholesteryl esters are removed from circulation via two pathways, either they are exchanged for triglycerides in apolipoprotein B containing lipoproteins, such as low-density lipoprotein (LDL), and subsequently delivered to the liver through the LDL receptor pathway or they are selectively extracted from mature HDL by scavenger receptor B1 for direct intrahepatic processing (7).

Because of the important role of HDL in RCT, synthetic HDLs (sHDL) are attractive candidates for the treatment of cardiovascular disease (8–11). Currently, two major sHDL nanomedicines, CER-001 and CSL112 that are pre-β HDL mimetics made with full-length apoA-I and different lipid compositions, have reached large-scale clinical trials. Yet, the results showed that weekly infusions of CER-001 or CSL112 in subjects with acute coronary syndrome (ACS) did not significantly reduce clinical cardiovascular events in comparison with placebo infusions (12,13). Although the primary end-point efficiency of CSL112 intervention was negative, there were preliminary indications of patients treated with CSL112 having numerically lower rates of cardiovascular death and myocardial infarction compared with placebo suggesting that some patient sub-groups under ACS could still benefit from this kind of treatment (14). Interestingly, a follow-up study reported that CSL112 infusions enhanced the cholesterol efflux capacity of plasma HDL fractions. It may be partially attributed to an increased rate of cholesterol esterification by LCAT that was shown to exhibit greater association with HDL fractions following infusion (15).

To date, no long-term, large-scale clinical trial involving sHDL particles has been conducted in the pre-ACS setting, largely due to the high production costs associated with apoA-I and the need for frequent intravenous administration over extended periods. A possible alternative to full-length apoA-I in sHDL infusion therapies is to use apoA-I mimetic peptides as they are cost-efficient, easily scalable and provide flexibility in modifying their size, physical properties and structure (16–18). The first apoA-I mimetic peptide, 18A, an 18-residue amphipathic helix analog, was designed to recreate apoA-I secondary structure and its ability to strongly bind lipids (19). Since then, several peptides have been developed from 18A sequence. It was shown that increasing the hydrophobicity of 18A enhance its ability to bind lipids and to promote cholesterol efflux (20). Longer bihelical peptides were designed by connecting two 18A peptides by a proline residues and show greater ability to efflux cholesterol from macrophages compared with the monomeric peptide (21,22). Another study discovered that designing bihelical peptides with helices of different hydrophobicity improve their specificity for promoting cholesterol efflux by the ABCA1 transporter (23).

The apoA-I mimetic peptide 22A (Esperion Therapeutics, ESP24218) was optimized to facilitate the activation of LCAT (24). It is the first apoA-I mimetic peptide to reach clinical trials as HDL-targeted therapy (ETC-642) in dyslipidemia patients (25). Results showed that the formulation was safe and well tolerated at the dose levels tested in the study and induced a rapid dose-related cholesterol mobilization. Despite these promising results, the development of ETC-642 was terminated in 2006 (26). Nevertheless, peptide-based sHDL particles continue to be investigated not only as a potential therapeutic agents for cardiovascular diseases but also as valuable tools for elucidating the mechanistic principles of various steps in RCT and their roles in atherosclerosis (27). Further functional studies are still needed to provide detailed insights into how the structure and composition of HDL particles affect their mechanisms of action in RCT, which includes the understanding of their interactions with macrophage lipid transporters, LCAT and endogenous HDL in plasma. Recent articles have demonstrated that sHDL particles composed of 22A peptides and DMPC lipids are remodeling endogenous HDL in human plasma, resulting in increased levels of lipid-poor HDL particles that possess the greatest capacity to absorb cellular cholesterol (28,29). Previous studies on different apoA-I mimetic peptides exhibited that upon remodeling the peptides specifically associate to endogenous HDL in plasma (30,31). Similar endogenous HDL remodeling has also been reported in the case of CSL112, composed of full length apoA-I and more physiological-like lipid composition, when injected into human (32). It has been argued that this remodeling capability of sHDL particles is the main reason behind the increased cholesterol efflux capacity of HDL particles in plasma after intravenous injection. Yet, there are no studies investigating how sHDL particles which are not remodeling endogenous HDL in plasma would perform in this context.

In this study, we used the subsequent developments in 18A peptide design to create new apoA-I mimetic peptides. We designed 22A-F, by adding a phenylalanine residue at the C-terminal of 22A, inducing an increase in hydrophobicity and 22A-P-18A, a bihelical peptide with helices of different hydrophobicity, by connecting 22A and 18A sequences by a proline residue. We hypothesized that sHDL particles made with these more hydrophobic and longer apoA-I mimetic peptides will better promote cholesterol efflux from macrophages *in vitro* and be more stable in human plasma compared to 22A-sHDL particles. In addition, we hypothesized that these modifications would retain the LCAT activation potential observed with the 22A peptide, as both newly designed peptides demonstrated the ability to interact with the peptide binding site of LCAT in our molecular dynamics (MD) simulations, a site that we have previously identified as critical for LCAT activation (33,34). We report the functional effects of these different peptides on the formation of sHDL, proteolytic stability, cholesterol efflux from macrophage cells and activation of LCAT *in vitro* as well as their ability to remodel endogenous HDL in human plasma.

## Material and methods

### Materials

ApoA-I mimetic peptides 22A (PVLDLFRELLNELLEALKQKLK), 22A-F (PVLDLFRELLNELLEALKQKLKF) and 22A-P-18A (PVLDLFRELLNELLEALKQKLKPDWLKAFYDKVAEKLKEAF) were synthesized by Peptide Protein Research Ltd. (Fareham, UK). 2-Dimyristoyl-sn-glycerol-3phosphocholine (DMPC) and ergosta-5,7,9,(11),22-tetraen-3β-ol (dehydroergosterol, DHE) were obtained from Avanti Polar Lipids Inc. (Alabaster, AL). Cholesterol oxidase, monoclonal anti-apoA-I antibody, formic acid and acylCoA-cholesterol acyltransferase (ACAT) inhibitor Sandoz 58-035 were obtained from Sigma-Aldrich (St. Louis, MO). Recombinant human lecithin cholesterol acyltransferase (LCAT) was synthesized by ProSpec-Tany TechnoGene Ltd. (East Brunswick, NJ). Polyacrylamide gels 4-20% and 0.2 µm nitrocellulose membrane were obtained from Bio-Rad (Hercules, CA). NativeMark HMW Protein Standards and goat anti-mouse IgG secondary antibody HRP were purchased from Thermo Fisher Scientific (Waltham, MA). Pooled human plasma from normal healthy donors was purchased from Innovative Research, Inc (Novi, MI). J774A.1 cells and Dulbecco’s Modified Eagle’s Medium (DMEM) were obtained from ATCC (Manassas, VA). ^3^H-cholesterol was bought from Revvity, Inc. (Waltham, MA). Iodoacetamide, 4-aminoatipyrine, phenol and peroxidase were purchased from Merck Life Science (Darmstadt, Germany).

### MD simulation

The detailed methods for preparation and analysis of MD simulations can be found in Supplemental Information. Coarse-grained molecular dynamics simulations using the MARTINI 3.0 force-field were run with the GROMACS software package (version 2020.2) (35). Initial coordinates for peptides were attained with PEPFOLD4 software (36) and for LCAT the protein preparation wizard of SCHRÖDINGER software package was used to prepare PDB ID 6MVD. For 22A, simulations were prepared by placing 200 DMPC and 28 peptides randomly into a cubic box. For the combined peptide mass to be approximately equal between systems, 27 peptides were placed to 22A-F systems and 15 peptides to 22A-P-18A systems. The random lipid-peptide complex was compressed into a flat plane and LCAT was placed on top of it. The system solvated with ∼ 63500 water beads. After energy minimization, the production systems were simulated until 32 µs with a 20 fs timestep and with coordinates without water written every 1 ns. All simulations were run as triplicates. Analyses were started at 2 µs, affording 90 µs of analysis per peptide in total. VMD program was used to generate snapshot images (37).

### sHDL preparation

DMPC phospholipids dissolved in chloroform were dried for 1 hour in a glass vial with Rotavapor R-200 (BUCHI, Switzerland). The resulting thin lipid layer was hydrated with 1/50X PBS pH 7.4 and homogenized by sonication in an ultrasonic bath at room temperature. Peptides (22A, 22A-F, 22A-P-18A), dissolved in 1/50X PBS pH 7.4, were added to reach a 2:1 w/w lipid to peptide ratio. Samples were cycled three times between 50°C (5 min) and ice (5 min) to facilitate sHDL formation. Samples were stored at +4°C until use.

### Dynamic light scattering

The size distribution profile of sHDL particles was measured by dynamic light scattering (DLS) with a Malvern Zetasizer APS at 25°C. The results are shown as volume distribution of hydrodynamic diameters. Data are represented as the average of three measurements.

### Size exclusion chromatography

Samples were analyzed on an NGC chromatography system (Bio-Rad) with a Superdex 200 Increase column (Cytiva) equilibrated in 1X PBS at room temperature. The flow rate was 0.4 mL/min and absorbance was measured at 190 nm.

### Circular dichroism

CD spectra of the free peptides and the peptides in sHDL particles were obtained using a J-815 circular dichroism spectropolarimeter (Jasco) equipped with a Peltier temperature controller. Samples were diluted to a final peptide concentration of 0.01 mg/mL in 1/50 X PBS buffer, pH 7.4. Data were acquired in a 1 cm quartz cuvette at 37°C with a data interval of 0.2 nm over a wavelength range of 200–260 nm and scan rate of 100 nm/min. Four spectra were recorded, averaged and buffer subtracted for each sample. The percentage of helix was estimated using the CDPro analysis software with the program CONTIN (38).

### Transmission electron microscopy

sHDL particles were imaged on a Hitachi HT7800 electron microscope at 100 kV. Samples were diluted to a final peptide concentration of 10 µg/mL in 20 mM HEPES, 120 mM NaCl, 1 mM EDTA buffer. 3 μL of sample was applied on an electron microscopy grid (200 mesh Cu) coated with carbon. After 45 second excess liquid was removed. The grid was washed three times in individual droplets of ultrapure water. The grid was then negatively stained with 2% uranyl acetate, first for 5 seconds and then for 60 seconds. Filter paper was used to blot the grid between the two staining steps. The grid was air-dried before measurement. The microscope was operated at a magnification of 40 000X, resulting in a pixel size of 2.8 Å.

### Cholesterol efflux assay

J774A.1 macrophages were cultured to 80% confluency in 48-well plates in DMEM with 10% fetal bovine serum and penicillin-streptomycin. Cells were then labeled for 24 hours with 1 µCi/mL of ^3^H-cholesterol in DMEM with 0.5% BSA and 5 µg/mL ACAT inhibitor Sandoz 58-035. After washing the cells with 1X PBS, sterile filtered sHDL particles were added at 3, 6, 10, 15 and 20 µM peptide concentration in DMEM with 0.5% BSA and 5 µg/mL ACAT inhibitor Sandoz 58-035. The DMPC control sample has been prepared as sHDL particles but without addition of peptide after sonication. Medium was collected after 18 hours of incubation and centrifuged (10 000 rpm, 10 min) to remove floating cells. Cells were rinsed with 1X PBS and lysed by addition of 0.1% SDS, 0.1 M NaOH buffer. Cell lysates were centrifuged (10 000 rpm, 15 min) after 30 min incubation at room temperature. Radioactive counts were measured by liquid scintillation counting in media and cell lysates. Cholesterol efflux was determined by calculating the ratio between media count and the sum of media and cell counts. The assay was performed in triplicate.

### LCAT cholesterol esterification assay

The fluorescent sterol DHE was used as the substrate in this assay to mimic the behavior of cholesterol (39). DHE-containing sHDL were prepared via the same method described in the sHDL preparation section. Samples were protected from light during the preparation and the assay as DHE is light sensitive. DMPC and DHE were combined at a 9:1 molar ratio in chloroform and dried with Rotavapor R-200 for 1 hour. The resulting thin layer was hydrated with 1/50 X PBS, 1mM EDTA pH 7.4 and sonicated for 1 hour. Peptides (22A, 22A-F, 22A-P-18A) were added to reach a 2:1 w/w lipid to peptide ratio. Samples were incubated for 5 min at 50°C followed by 5 minutes on ice. This heating-cooling cycle was repeated 3 times. Recombinant LCAT was incubated with DHE-sHDL at different DHE concentration (0, 4, 8, 12, 24 and 36 µM) in assay buffer (1/50X PBS, 1 mM EDTA, 60 µM albumin, pH 7.4) for 1 hour at 37°C. The final recombinant LCAT concentration was 2.5 µg/mL. Reactions were stopped by adding stop solution (5 U/mL cholesterol oxidase, 1/50X PBS, 1 mM EDTA, 7% Triton X-100, pH 7.4) and incubated for 1 hour at 37°C to quench the fluorescence of unesterified DHE. The DHE-ester fluorescence was detected using Varioskan LUX at an excitation wavelength of 325 nm and an emission wavelength of 425 nm. A standard curve was obtained by measuring the fluorescence of DHE-sHDL at different concentrations. All fluorescence data were divided by the slope of the standard curve to obtain the concentration of DHE ester in the reaction. The DHE esterification rate (concentration of DHE ester divided by the time of the assay) was plotted against the concentration of DHE and fitted to a Michaelis-Menten kinetic equation to determine the kinetic parameters V_max_ and K_m_. The assay was performed in triplicate.

### Remodeling of HDL by free peptide or sHDL in human plasma

Human plasma was incubated with 1/50X PBS (control), free peptides and sHDL particles at a final peptide concentration of 1 mg/mL for 1 hour at 37°C. The reaction was quenched by 10-fold dilution in 50% sucrose solution. 10 µL of the diluted samples and 10 µL of the NativeMark HMW protein standards were loaded into a 4-20 % polyacrylamide gel equilibrated in 25 mM Tris, 1.92 M glycine, pH 8.3 buffer. The gel was run for 2.5 hours at 200 V. The ladder line was cut from the gel and analyzed via Coomassie staining. The rest of the gel was transferred onto a 0.2 µm nitrocellulose membrane in Bio-Rad’s Trans-Blot Turbo System for 10 minutes at 1.3 A. The membrane was blocked for 1 hour with 5% nonfat dry milk in TBST (20 mM Tris, 150 mM NaCl, pH 7.6, 0.1 % tween-20). The membrane was incubated overnight at 4°C in 5% nonfat dry milk TBST containing a 1:3000 dilution of a monoclonal anti-apoA-I antibody produced in mouse (stock concentration 0.2 mg/mL). After extensive wash in TBST, the membrane was incubated for 1.5 hours in 5% milk TBST containing a 1:7000 dilution of a goat anti-mouse IgG secondary antibody HRP (stock concentration 1.5 mg/mL). After extensive wash in TBST, the membrane was incubated for 5 minutes with an enhanced chemiluminescence reagent. The membrane was imaged by chemiluminescence detection using the ChemiDoc MP imaging system (Bio-Rad). The standards were: ferritin (480 kDa, 12.2 nm), B-phycoerythrin (242 kDa, 10.1 nm), lactase dehydrogenase (146 kDa, 8.1 nm) and BSA (66 kDa, 7.1 nm) (40,41). We have tested that the anti-apoA-I antibody does not recognize 22A, 22AF and 22A-P-18A. The assay was performed in duplicate.

### LCAT cholesterol esterification in human plasma

Cholesterol esterification rate was determined by a colorimetric assay as previously described (42,43). Human plasma was incubated with 1/50X PBS (control), free peptides and sHDL particles at a final peptide concentration of 1 mg/mL for 1 hour at 37°C. Reactions were stopped immediately after the addition of peptides or sHDL particles to plasma and after 1 hour incubation by addition of 0.15 M iodoacetamide and placing the tubes on ice. The concentration of red quinone produced in the reaction in presence of 0.7 U/mL cholesterol oxidase, 0.7 U/mL peroxidase 15 mM phenol and 1 mM 4-aminoantipyrine concentration was photometrically measured with a Cary 100 UV-visible spectrophotometer (Agilent Technologies) at 510 nm. A standard curve was obtained by performing the same reaction at different cholesterol concentrations. Cholesterol esterification rate was calculated by subtracting the concentration of unesterified cholesterol at the beginning of the reaction and after 1 hour incubation. The assay was performed in triplicate.

### Stability of peptide in human plasma

22A, 22A-F and 22A-P-18A peptides were incubated at a final peptide concentration of 0.25 mg/mL with human plasma at 37°C, either in their free form or in the sHDL formulations. Aliquots were removed immediately after the addition of peptide to plasma and after 24 hours incubation. Plasma proteins were precipitated by the addition of two volumes of acetonitrile/ethanol (1:1 v/v), centrifuged at 10000 rpm for 10 minutes and the supernatants were transferred to low-binding vial inserts. The proportion of intact peptide in the sample was determined by liquid chromatography-high-resolution mass spectrometry (LC-HRMS), as described in detail in the Supplemental Information. Briefly, samples were analyzed using a Waters Xevo QTOF high-resolution mass spectrometer coupled to a Waters Acquity UPLC system. Chromatography was performed using a C_18_ column (Waters Acquity UPLC BEH C-18, 1.7 µm, 100 mm × 2.1 mm) with a 2 µL injection volume. The mobile phase consisted of (A) water and (B) methanol, both containing 0.1% formic acid. An LC gradient was run at a flow rate of 0.3 mL/min. Mass spectra were acquired using an electrospray ionization source with a 1.5 kV capillary voltage in positive ion mode, with a scan range of m/z 120–1500. The assay was performed in triplicate.

### Statistics

Data are represented as mean ± standard deviation (SD) of triplicate measurements. Statistical analysis was performed with a one-way analysis of variance (ANOVA) followed by a two-tailed Dunnett’s post-hoc test, using IBM SPSS Statistics software.

## Results

### Design of new amphiphilic apoA-I mimetic peptides

The apoA-I mimetic peptides used for this study are described in Figure 1. Two new apoA-I mimetic peptides were designed by modifying the sequence of the well-studied 22A peptide. The peptide 22A-F has one extra phenylalanine residue at the C-terminal of 22A. The peptide 22A-P-18A is a bihelical peptide containing the 22A and 18A amphipathic helices linked by a proline residue. We selected the C-terminal end of 22A for insertion of the modifications because our previous simulations studies (33,34) showed that the N-terminal end of 22A becomes buried when it interacts with LCAT. Helical-wheel diagrams illustrate the amphiphilic nature of the peptides, as hydrophobic amino acids are clustered on one side of the helices (Fig. 1A). 22A-F peptide has a slightly higher affinity to hydrophobic surface than 22A and 22A-P-18A peptides, as assessed by their retention time on a C_18_ reverse-phase UPLC column (6.24 min, 6.14 min and 6.06 min, respectively) (Table S3). Coarse-grained molecular dynamics simulations were employed to investigate the location and orientation of apoA-I mimetic peptides in sHDL particles and to verify their accessibility to the peptide binding site of LCAT. In accordance with our previous results (33,34), peptides and DMPC phospholipids spontaneously self-assembled into nanodiscs structures that are stable throughout the duration of the simulations. Peptides are located on the edge of the discs to shield the hydrophobic acyl chains of the phospholipids from the water phase. They are mostly assembled in a double belt-like arrangement as in it the case for apoA-I in HDL particles (44,45). For most of 22A-P-18A peptides, we observed the formation of a turn between the two amphipathic helical segments. In all simulations, LCAT moved to the perimeter of the disc and all the peptides utilized the peptide binding site for binding LCAT (Fig. 1B). On average, 22A bound to the peptide binding site of LCAT more frequently than 22A-F and 22A-P-18A (34.2 ± 9.4 %, 25.8 ± 2.5 % and 13.9 ± 1.6 %. respectively) (Fig. S1A). Analysis of the spatial position of the peptides with respect to the lipids in the sHDL particles revealed that in the 22A-P-18A-sHDL simulations, the 22A fragments of the 22A-P-18A peptides are steered to the rim of the disc more strongly than in the 22A-sHDL simulations where 22A peptides are also occasionally located closer to the centre of the disc (Fig. 1C). Further analysis of the MD simulations showed that both 22A-F and 22A-P-18A had slower rotation on the phospholipid bilayer than 22A (Fig. S1B). Unlike 22A and 22A-F, which primarily utilized early residues of the 22A fragment to bind to LCAT, 22A-P-18A utilized early residues of the 18A fragment (Fig. S2 and Table S1).

**Figure 1:**
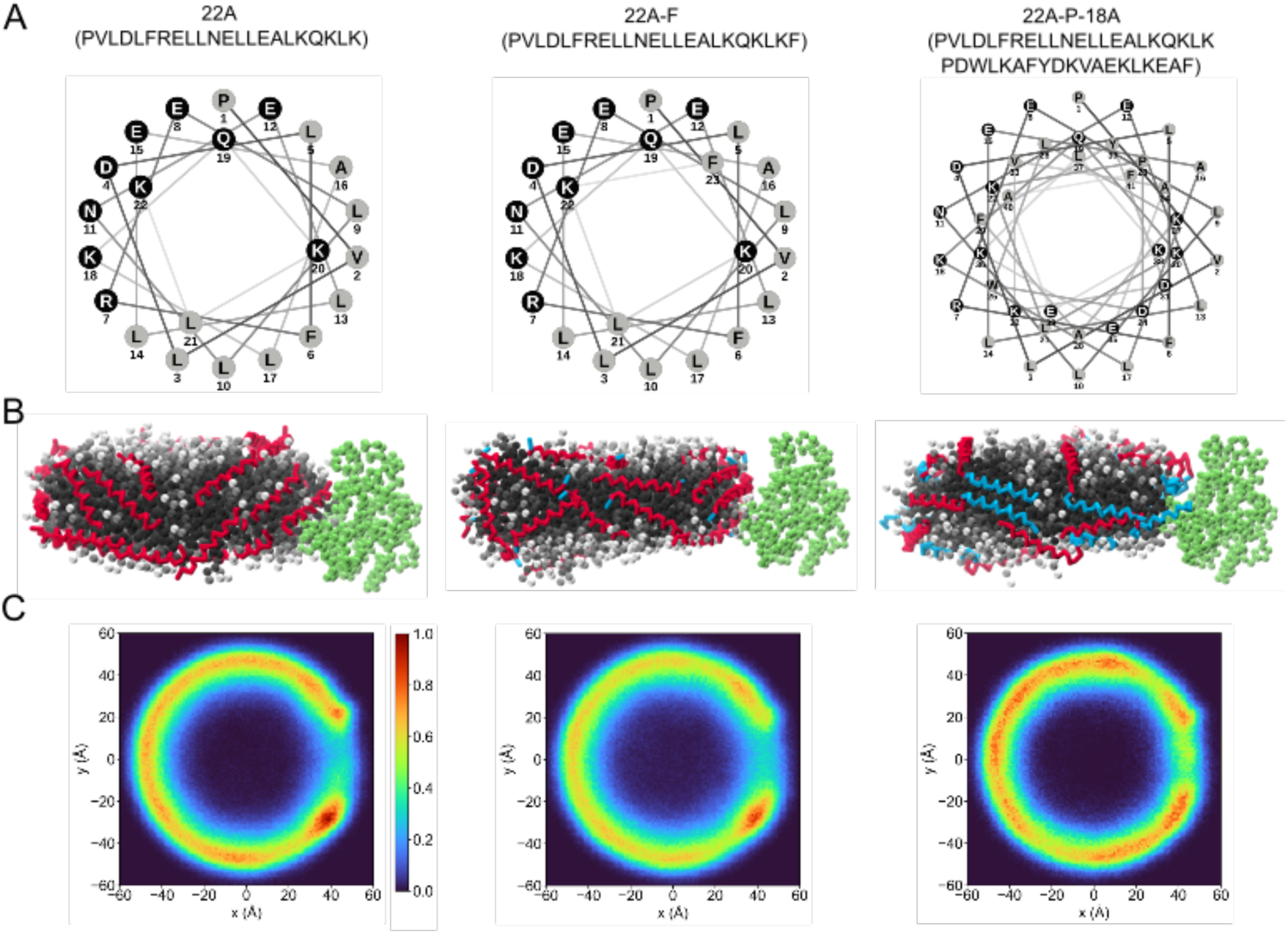
Design of amphiphilic peptides by modifying the 22A apoA-I mimetic peptide sequence. (A) Helical wheel plot generated using the online NetWheels server of the peptides used in this study and their sequence. Hydrophobic amino acids are color coded in gray. (B) Molecular dynamics simulation snapshots of sHDL particles showing their molecular organization. DMPC and LCAT are rendered as spheres and peptides as sticks. DMPC is colored in greyscale, peptides in red and blue, with the 22A sequence in red, and LCAT in green. Water beads are removed from the figure for clarity. (C) Peptide position heatmaps in sHDL particles, top view. The colorbar was normalized by the maximum value of all systems. The coordinates were transformed based on the center of mass of the sHDL particle, the sHDL particle normal, and the center of mass of LCAT, so that LCAT is always at y = 0 and positive x. The volume taken by LCAT translates into a gap in the heatmap.

### Preparation and biophysical characterization of lipid-free peptides and sHDL particles

We first tested the ability of the peptides to form homogeneous sHDL particles by combining them with DMPC lipids at a 2:1 lipid:peptide w/w ratio. All three peptides were able to form homogeneous sHDL particles with an average hydrodynamic diameter ranging from 8.9 to 10.1 nm as determined by DLS (Fig. 2A and Table 1). Size exclusion chromatography was used to characterize the purity of sHDL particles. No peak was observed at the retention time of lipid-free peptide (40.3 min) indicating that all peptides were associated with lipids (Fig. 2B). This is consistent with our MD simulation results, which show that all peptides interacted strongly with lipids, with no peptides observed to dissociate or remain unbound during the simulations. The helical content of peptide in lipid-free form and in sHDL particles was determined by circular dichroism. The three peptides have roughly 80 % of helical content in both lipid-free form and sHDL particles (Fig. 2C and Table 1). This indicates that all peptides form stable amphiphilic α-helices, and their interaction with lipids does not alter their secondary structure. All sHDL particles have discoidal morphology, as determined by TEM (Fig. 2D).

**Figure 2:**
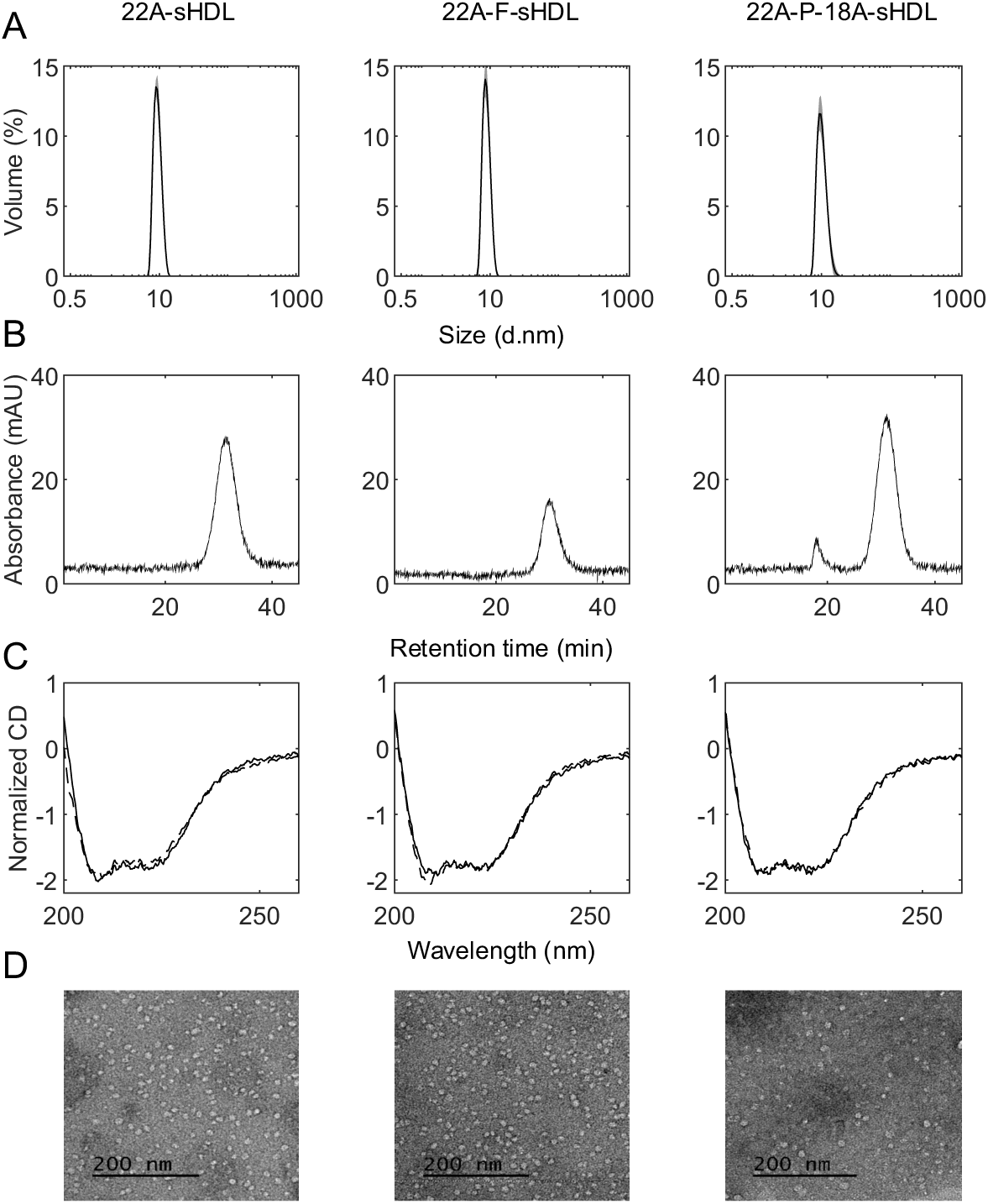
Physicochemical characterization of sHDL particles prepared with DMPC and different peptides. (A) Volume based size distribution determined by dynamic light scattering. Data are represented as mean ± SD of triplicate measurements. (B) Size exclusion chromatography, absorbance was measured at 190 nm. (C) Circular dichroism spectra of lipid-free peptides (dashed lines) and sHDL particles (solid lines). (D) Negative-stain TEM images.

**Table 1:** Physicochemical parameters of sHDL particles prepared with DMPC and different peptides. Circular dichroism parameters were calculated using CDPro analysis software. Data are presented as mean ± SD.

|  |  | 22A | 22A-F | 22A-P-18A |
| --- | --- | --- | --- | --- |
| Size measured by DLS (nm) |  | 9.4 ± 1.3 | 8.9 ± 1.1 | 10.1 ± 1.6 |
| CD parameter (% helix) | Lipid-free peptide | 79.7 ± 1.3 | 82.4 ± 1.4 | 80.0 ± 1.3 |
|  | sHDL | 83.4 ± 1.4 | 80.0 ± 1.4 | 78.2 ± 1.4 |

### Effects on macrophage cholesterol efflux

During RCT, HDL particles promote net cholesterol efflux from macrophage foam cells which is beneficial in reducing lipid accumulation in atherosclerotic plaques. To test the ability of sHDL particles made with different peptides to stimulate cholesterol efflux, J774A.1 macrophages labeled with ^3^H-cholesterol were incubated with various concentrations of sHDL for 18 hours. All three sHDL formulations showed a concentration-dependent cholesterol efflux (Fig. 3 and Table S2). 22A-P-18A-sHDL induced increased cholesterol efflux in comparison with 22A-sHDL at all concentrations. At sHDL concentration of 10 µM and higher, 22A-F-sHDL promoted cholesterol efflux more efficiently than 22A-sHDL. Our results show that either adding a single phenylalanine to the C-terminal end of 22A or linking it to the more lipophilic peptide 18A similarly enhances cholesterol efflux. Additionally, the DMPC control formulation exhibited cholesterol efflux from macrophages, although lower in magnitude when compared with sHDL particles. This likely reflects a passive non-ABCA1 mediated cholesterol efflux from the cells.

**Figure 3:**
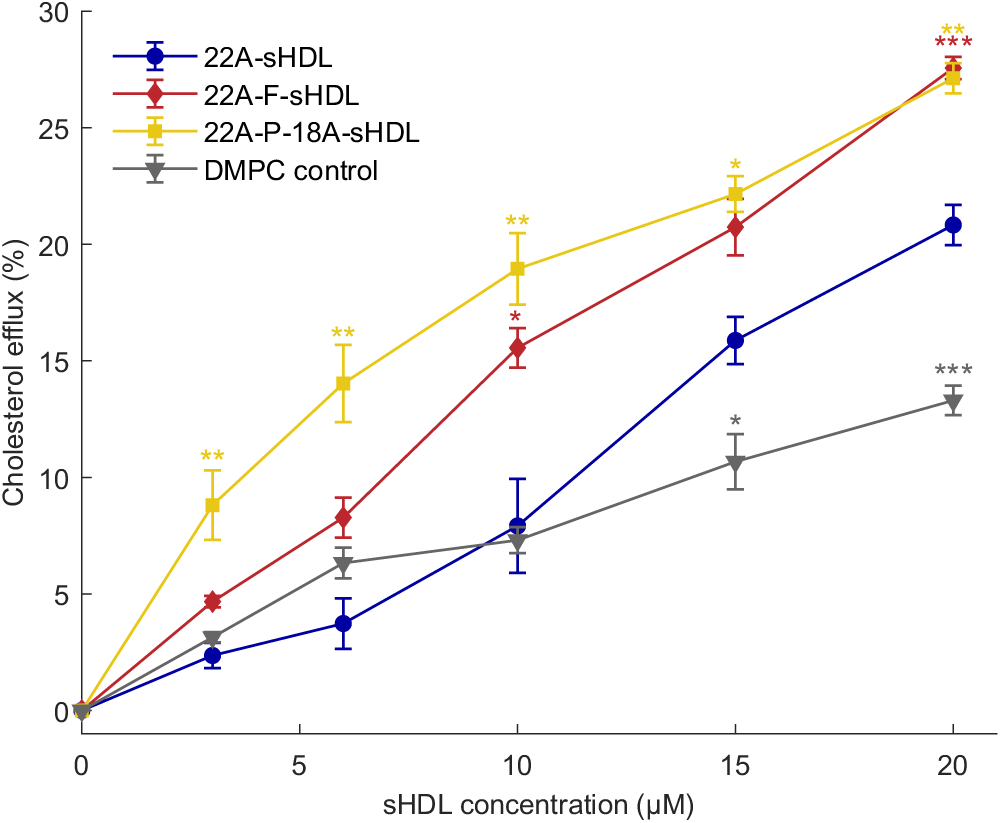
Cholesterol efflux from J774A.1 macrophage cells to sHDL particles. Cholesterol efflux was measured as the percentage of ^3^H-cholesterol in the medium after 18 hours of incubation with sHDL particles. Data are represented as mean ± SD of triplicate measurements. Statistical significances were compared with 22A-sHDL with one-way ANOVA and Dunnett’s post-hoc test. Each concentration point was treated independently. *P < 0.05, **P < 0.01, ***P < 0.001.

### Effects of sHDL interaction with recombinant LCAT

The interaction in plasma between HDL particles and the LCAT enzyme is an essential step of RCT as it promotes maturation of HDL particles via cholesterol esterification. To assess the impact of different peptides in sHDL formulation on LCAT acyltransferase activity, fluorescent cholesterol-like dehydroergosterol (DHE) was incorporated to sHDL particles and used as a substrate. The rate of DHE esterification by recombinant LCAT in sHDL particles was measured at different DHE concentrations and fitted to the Michaelis-Menten equation to determine the kinetic parameters K_m_ and V_max_ (Fig. 4A and Table 2). The catalytic rate constant K_cat_ was used to characterize the efficiency of LCAT esterification of DHE-containing sHDL particles (Fig. 4B and Table S2). All three sHDL formulations were able to activate LCAT. While 22A-F-sHDL did not have a significant difference in catalytic rate constant compared to 22A-sHDL (K_cat_ = 47.6 ± 1.4 and 46.1 ± 0.8 h^-1^, respectively), 22A-P-18A-sHDL significantly improved LCAT catalytic efficiency by 35 % (K_cat_ = 62.2 ± 6.2 h^-1^) in comparison with 22A-sHDL. Thus, our results indicate that adding a phenylalanine or the 18A peptide to the C-terminal end of 22A does not disrupt the peptides’ ability to activate LCAT and, surprisingly, even enhanced it in the case of 22A-P-18A.

**Figure 4:**
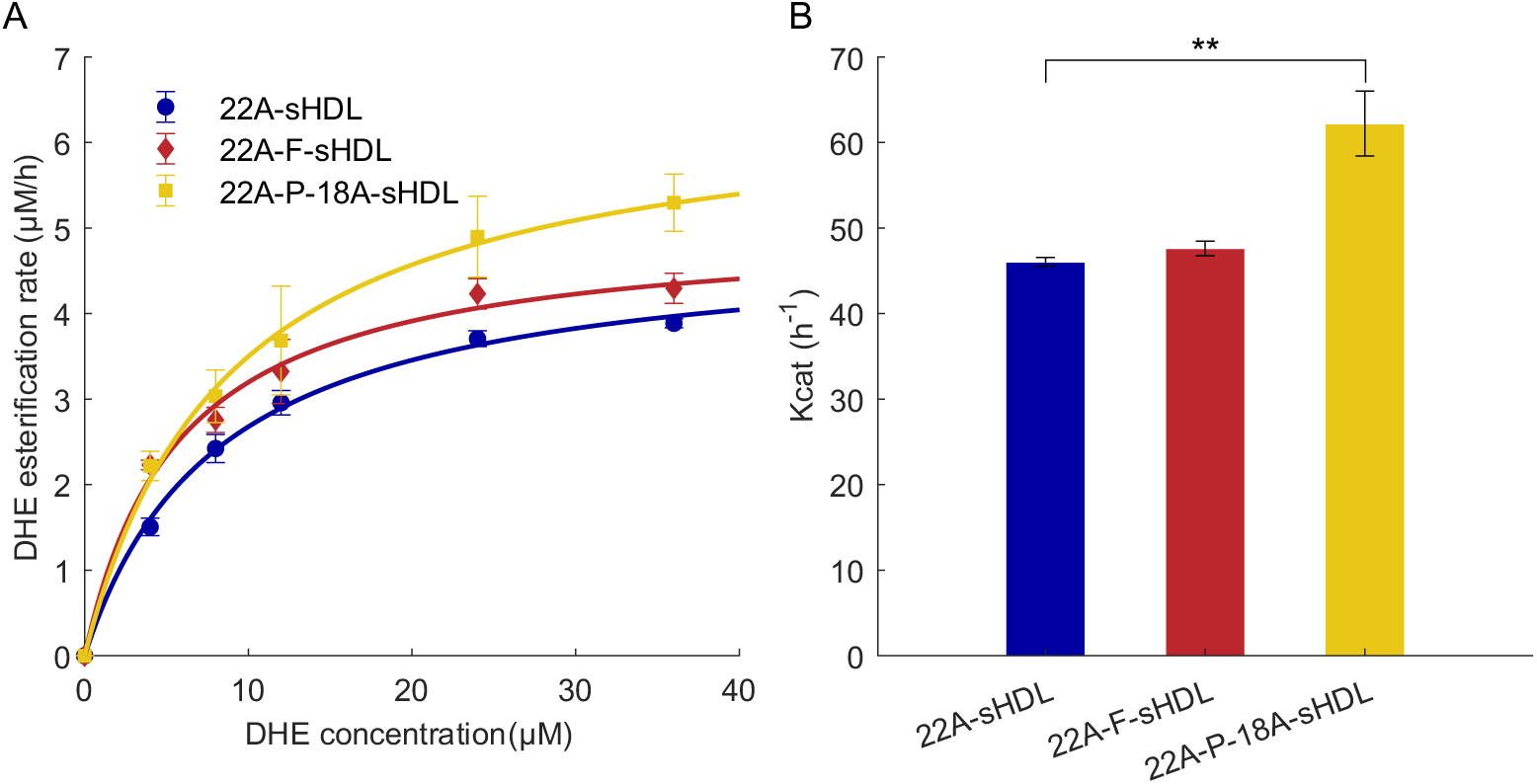
Effect of peptide in sHDL particles on the esterification rate of recombinant LCAT. (A) DHE esterification catalyzed by recombinant LCAT of different DHE-containing sHDL particles. Data were fitted to the Michaelis-Menten kinetic equation to determine the kinetic parameters V_max_ and K_m_. (B) Catalytic efficiency of LCAT esterification of different DHE-containing sHDL particles. K_cat_ is calculated as V_max_ divided by the enzyme concentration. Data are represented as mean ± SD of triplicate measurements. Statistical significances were compared with 22A-sHDL with one-way ANOVA and Dunnett’s post-hoc test. *P < 0.05, **P < 0.01.

**Table 2:**
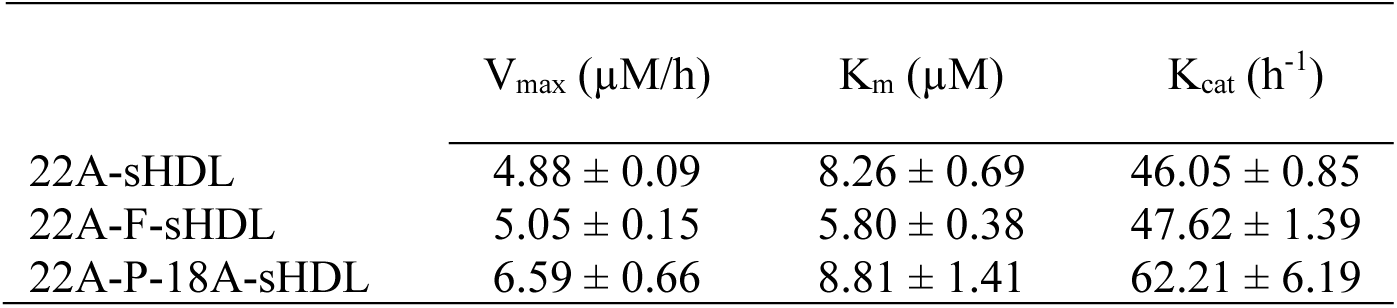
Kinetic parameters of the enzymatic reaction between recombinant LCAT and DHE-containing sHDL particles. Data are represented as mean ± SD of triplicate measurements.

### Effects on human plasma HDL remodeling

During the multiple steps of reverse cholesterol transport, HDL particles are present in distinct sizes and shapes (46). Lipid-poor apoA-I are small and dense nanoparticles with the greatest ability to induce phospholipid and cholesterol efflux from macrophages. The absorption of phospholipid and cholesterol leads to the formation of discoidal pre-β HDL particles. LCAT-mediated esterification of cholesterol generates mature, larger spherical α-HDL particles that are the most abundant HDL particles in human plasma. Previous studies have indicated that sHDL particles containing full length apoA-I or its mimetics remodel endogenous α-HDL particles, presumably through particle fusion, which results in temporal increase of discoidal pre-β HDL and lipid-poor apoA-I particles (31,32). Therefore, we investigated the ability of lipid-free peptides or sHDL particles to remodel endogenous lipoproteins. This was carried out by incubating human plasma with different formulations at a final peptide concentration of 1 mg/mL for 1 hour at 37°C. The subfractions of endogenous HDL were separated via native polyacrylamide gel electrophoresis and visualized by immunoblotting for apoA-I. The plasma sample treated with 1/50X PBS as a control contained mainly mature α-HDL particles. All lipid-free peptides and sHDL particles induced the remodeling of endogenous HDL, as we observe decreased levels of mature α-HDL, a slight increase in pre-β HDL levels and a major increase of lipid-poor apoA-I (Fig. 5 and Fig. S3). Our findings highlight that increasing the hydrophobicity of the 22A peptide, either by adding a phenylalanine residue or by linking it to the more lipophilic peptide 18A, does not prevent the remodeling of endogenous HDL in human plasma. However, this assay cannot rule out the presence of intact sHDL particles, as it only detects HDL fractions containing apoA-I.

**Figure 5:**
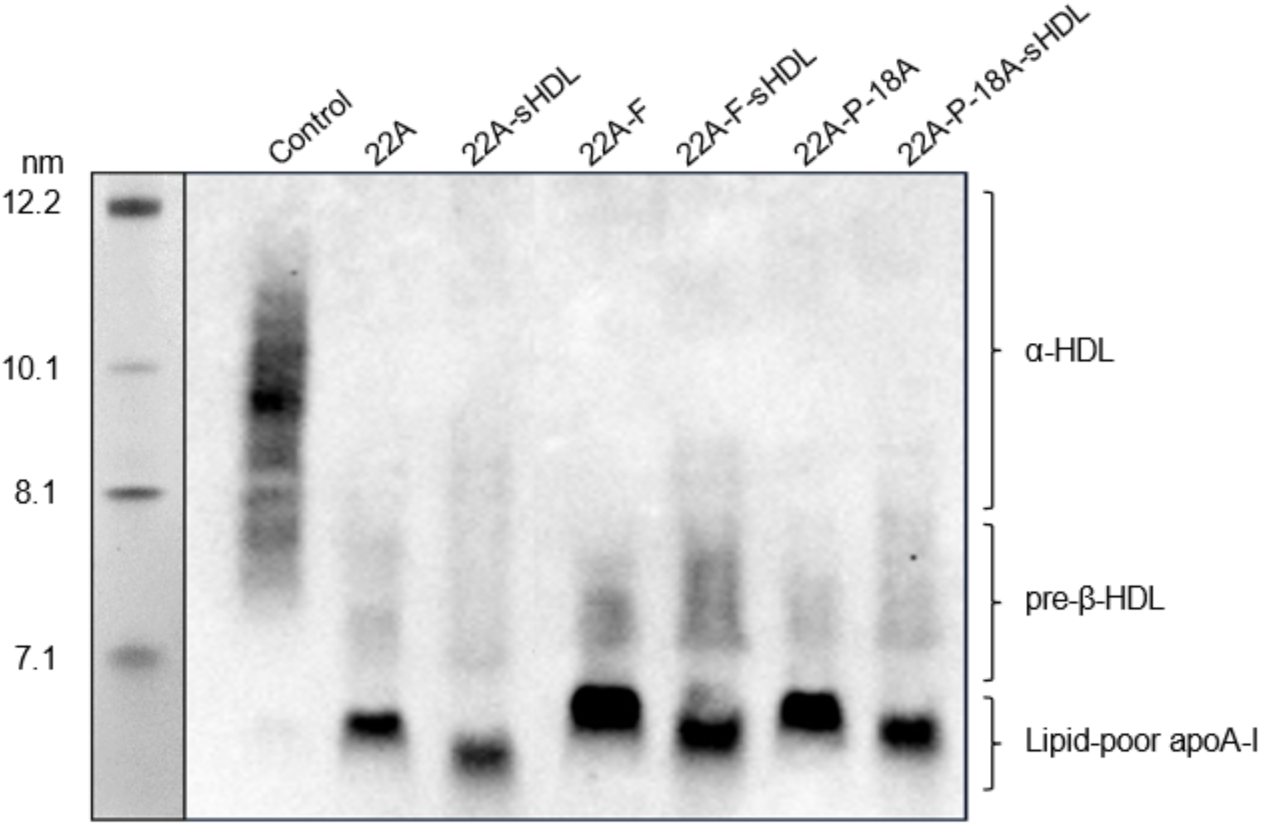
Remodeling of endogenous HDL in human plasma. Plasma samples were incubated with different lipid-free peptides and sHDL particles for 1 hour at 37°C. Lipoproteins were separated by nondenaturing polyacrylamide gradient gel electrophoresis followed by Western blotting with an anti-apoA-I antibody.

### Effects on LCAT activity in human plasma

As the recombinant LCAT activity assay employs DHE as a substrate and isolates the effects of 22A and its derivatives from native apoA-I, other apolipoproteins and endogenous HDL remodeling effects, we examined how the observed LCAT activities translate to a more physiological setting. Thus, the ability of endogenous LCAT to esterify cholesterol in human plasma in the presence of lipid-free peptides or sHDL particles was evaluated. The plasma cholesterol esterification rate is measured to quantify LCAT activity as this enzyme is predominantly responsible for the conversion of unesterified cholesterol to cholesteryl ester in human plasma. The changes in unesterified cholesterol after 1 hour incubation at 37°C with lipid-free peptides or sHDL particles at 1 mg/mL concentration were recorded and compared to the control, which corresponds to the basal activity of LCAT in human plasma (Fig. 6 and Table S4). All peptides were able to increase LCAT activity when formulated as sHDL particles but only 22A-P-18A sHDL particles are significantly enhancing LCAT activity in comparison with the control sample. Incubating human plasma with lipid-free peptides or with DMPC phospholipids (data not shown here) does not affect LCAT activity. Our results demonstrate that even though sHDL particles remodel the α-HDL fraction in human plasma, leading to the formation of pre-β HDL particles and lipid-poor apoA-I, the enhanced LCAT activation potency of 22A-P-18A is still preserved. Moreover, LCAT activity is higher for all sHDL formulations compared to the lipid-free peptides, which exhibit only basal activity levels. This indicates that the increased LCAT activation observed with sHDL formulations cannot be due to an increase of the pre-β HDL fraction.

**Figure 6:**
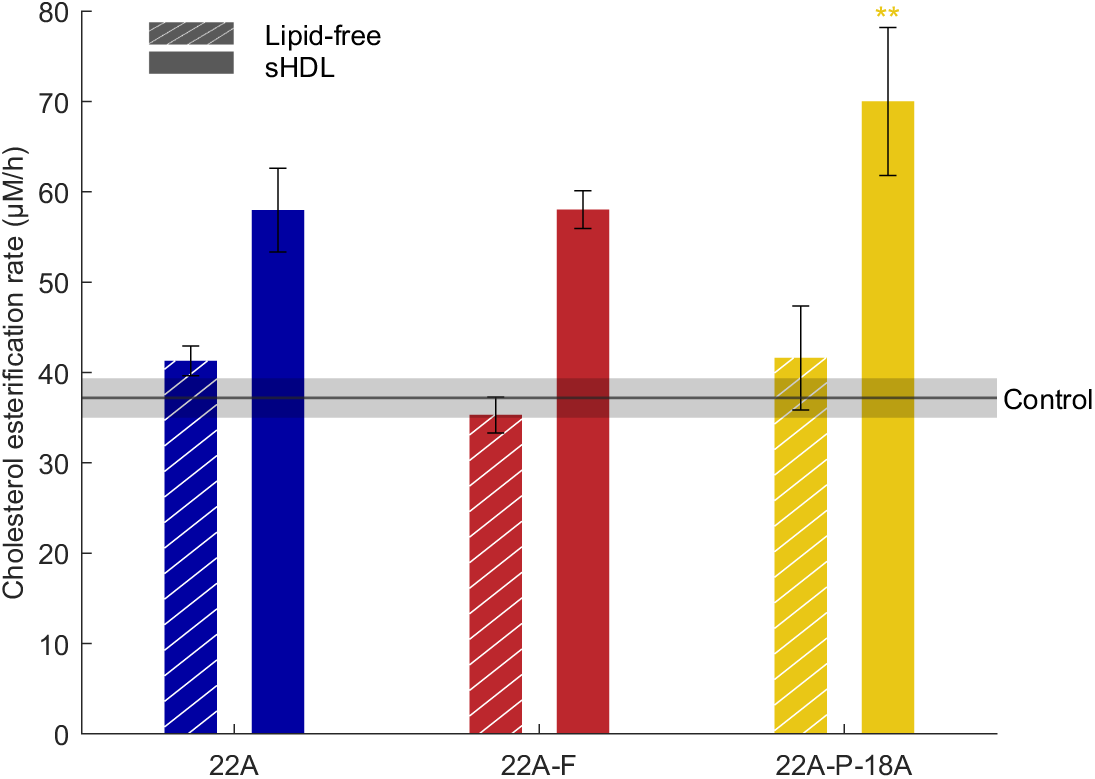
LCAT activation in human plasma. Peptides in their lipid-free from or in sHDL particles were incubated at 37°C for 1 hour with human plasma. The cholesterol esterification rate is reported. Data are represented as mean ± SD of triplicate measurements. Statistical significances were compared with the control (basal activity of LCAT in plasma) with one-way ANOVA and Dunnett’s post-hoc test. *P < 0.05, **P < 0.01, ***P < 0.001.

### Peptide proteolytic stability

Clinical use of peptides is often prevented by their poor proteolytic stability and short half-lives, as they experience enzymatic degradation by abundant proteases present in plasma (47). The proteolytic stability of 22A, 22A-F and 22A-P-18A peptides was investigated in their lipid-free form and in the sHDL particles. We monitored the disappearance of the intact peptides after 24 hours incubation in human plasma at 37°C by LC-HRMS (Fig. 7 and Table S3). We found that 22A and 22A-F were subject to minimal degradation during the 24 hours incubation, both in their lipid-free form and in the sHDL particles, with more than 95 % of intact peptide remaining. 22A-P-18A was overall less stable than 22A but it was more stable when formulated in sHDL particles compared to its lipid-free form, with 85.0 ± 5.2 % and 73.6 ± 6.3 % of intact peptide remaining respectively.

**Figure 7:**
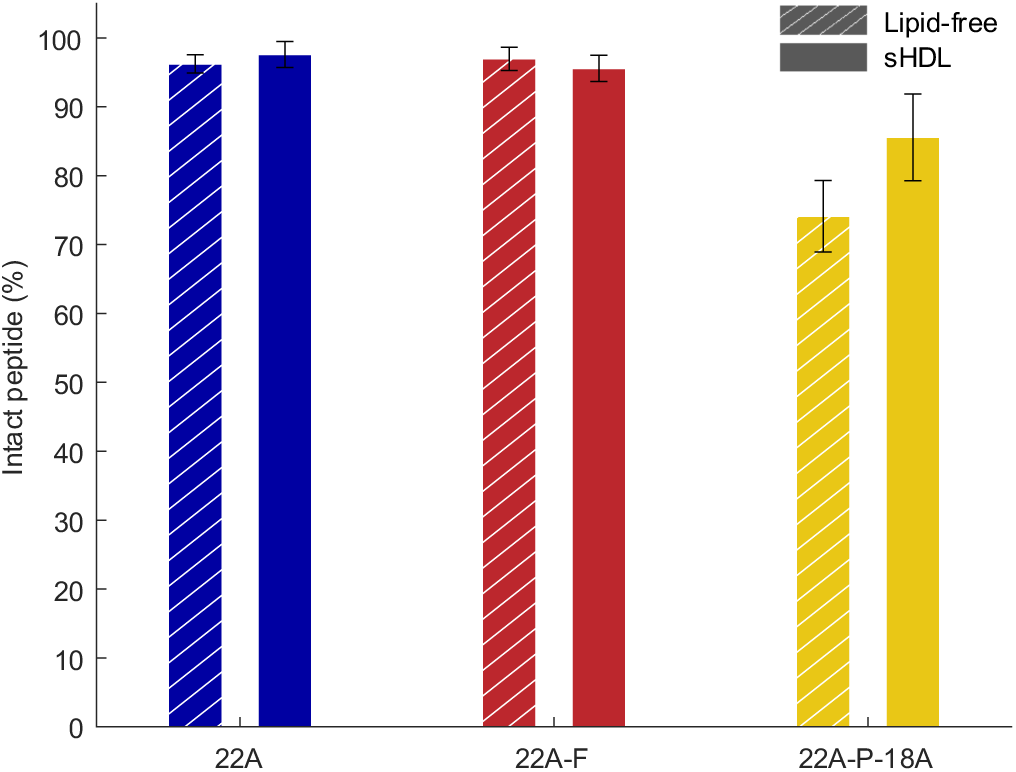
Susceptibility of peptides to proteolysis *in vitro*. Peptides in their lipid-free from or in sHDL particles were incubated at 37°C for 24 hours with human plasma. The percentage of intact peptide was monitored by LC-HRMS. Data are represented as mean ± SD of triplicate measurements.

## Discussion

In this study, we have developed two novel apoA-I mimetic peptides, 22A-F and 22A-P-18A, as derivatives of the well-characterized peptide 22A. 22A-F is more hydrophobic than 22A and 22A-P-18A is a bihelical peptide, longer than 22A and with helices of different hydrophobicity. We hypothesized that with these modifications we can improve the anti-atherosclerotic RCT properties of sHDL particles, specifically cholesterol efflux and plasma stability, while preserving LCAT activity at the same level compared to 22A. We systematically investigated the potency of the apoA-I mimetics peptides for cellular cholesterol efflux and LCAT activation *in vitro*, HDL remodeling in plasma and proteolytic stability.

The two novel peptides, in combination with DMPC phospholipids, could successfully form homogeneous, nanodisc shaped sHDL particles. DMPC phospholipids were used as they form fluid bilayers at physiological temperature and are known to facilitate LCAT binding with sHDL particles *in vitro,* although they have relatively short *in vivo* circulation time (28). MD simulation results suggest that the peptides are arranged in a double belt-like arrangement, like apoA-I in HDL particles, which is also consistent with a previous report on 18A-sHDL particles (48). Simulations further revealed that all peptides strongly interacted with lipids and no dissociation of peptides were detected. How the peptides interacted with LCAT is consistent with our previous modeling results (33,34).

We found that 22A-F and 22A-P-18A exhibited higher cholesterol efflux capacity to macrophages than 22A. This is consistent with previous studies showing that increasing peptide hydrophobicity enhances their cholesterol efflux functionality (49) and that bihelical peptides better stimulate cellular cholesterol efflux compared to monomeric peptides (22,31). This is likely because 22A-F-sHDL and 22A-P-18A-sHDL particles are better able to interact with cellular transporters than 22A-sHDL particles. However, the specificity of these sHDL particles for promoting cholesterol efflux by the ABCA1 transporter is not known. This is an important aspect as non-ABCA1-dependent cholesterol efflux has been associated with cytotoxicity (23). These non-specific interactions depend on the lipid affinity of the apoA-I mimetic peptides (50) while interactions with the ABCA1 transporter also require a cluster of negatively charged amino acids on the polar face of amphipathic helices (51). Both 22A and 18A peptides have glutamic acid and aspartic acid residues on the polar face of their helices (Fig. 1A) and it has been show previously that cholesterol efflux to 22A-sHDL and 18A-sHDL particles was primarily mediated by ABCA1 (29). Thus, it remains to be tested whether 22A-F-sHDL and 22A-P-18A-sHDL particles are efficient acceptors for ABCA1-mediated cholesterol efflux, but it seems highly probable because they have the necessary structural elements to mediate ABCA1 cholesterol efflux.

Our data indicate that 22A, 22A-F and 22A-P-18A peptides, in their lipid-free form or in sHDL particles, promote HDL remodeling in plasma to produce lipid-poor apoA-I and pre-β HDL particles. These particles have the greatest capacity to induce cellular cholesterol efflux and to enhance RCT. Thus, the observed elevation of HDL-cholesterol in patients after administration of 22A-sHDL particles (ETC 642) (25) could be explained by the ability of 22A to remodel HDL in plasma. Similarly, previous works suggest that the remodeling of HDL particles mediated by CSL112 may account for the elevation of cholesterol efflux capacity in plasma (15,32). While we cannot determine from our data how mechanistically sHDL particles interact and remodel endogenous HDL particles, other studies tracked the interactions of peptides with native HDL in human plasma *in vitro* (30,31). It was shown that peptides built as multimeric amphiphilic α-helices are binding to lipoproteins more selectively and with a higher affinity than monomeric peptides. Considering these results, it is most likely that the bihelical 22A-P-18A peptide will more effectively associate with lipoproteins in plasma than 22A and 22A-F peptides.

The ability of apoA-I mimetic peptides to activate LCAT was investigated using a combination of MD simulations and biomedical studies. In silico results highlight that all three peptides were able to interact with the peptide binding site of LCAT that was previously shown to be the principal binding site for the peptides (33,34). Our experiments showed that 22A-P-18A sHDL particles were more effective in promoting cholesterol esterification mediated by recombinant LCAT than 22A-sHDL particles. Comparably, only 22A-P-18A sHDL particles were able to significantly increase endogenous LCAT activity in human plasma. Interestingly, endogenous HDL particles are remodeled similarly by the lipid-free peptides and the sHDL particles but lipid-free peptides do not affect the cholesterol esterification rate in human plasma. Our hypothesis is that, although endogenous HDL are remodeled, there are still intact sHDL particles present in plasma that are better LCAT activators when compared to pre-β HDL particles. If this is the case, the intact sHDL particles are able to acquire unesterified cholesterol and interact with endogenous LCAT to promote cholesterol esterification.

A previous study reported that 22A is rapidly hydrolyzed in rat plasma and lacks its C-terminal lysine (28), whereas we observed high proteolytic stability of 22A and 22A-F both in their lipid-free form and in the sHDL particles when incubated with human plasma. This difference can be explained by the source of plasma used, as it was shown that plasma from human sources seems to degrade peptides less extensively that rat analogues (52). 22A-P-18A has lower proteolytic stability that 22A, but it was still quite resistant to degradation in the sHDL particles, with only 15 % lost after 24h incubation with human plasma. Furthermore, 22A-P-18A may have a longer half-life *in vivo* relative to 22A since peptides with higher molecular weight are more resistant to renal clearance (53). This is very promising as a short circulatory half-life *in vivo* is a common limitation of peptide-based therapeutics.

To conclude, we successfully designed new apoA-I mimetic peptides, 22A-F and 22A-P-18A, that are functionally superior to the 22A peptide. They both demonstrated higher cholesterol efflux and, in the case of 22A-P-18A, superior ability to activate LCAT *in vitro*. In addition, they exhibited relatively good proteolytic stability *in vitro* in the sHDL formulation. Although these modifications did not affect the ability of sHDL particles to remodel endogenous HDL fractions in human plasma, the LCAT activation assay conducted in human plasma suggest that intact sHDL particles may still be present. The proportion of these intact particles, which might depend on their specific peptide and lipid compositions, is highly relevant in the context of sHDL-mediated drug delivery and should be investigated more closely in the future. Nevertheless, *in vitro* trends of different peptide-based sHDL formulations may not translate to anti-atherosclerotic effects *in vivo*, as demonstrated in several studies (28,29,54). Therefore, the impact of these novel peptides on RCT and atherosclerosis progression should be evaluated using appropriate *in vivo* models.

## Acknowledgment

The authors acknowledge the CSC−IT Center for Science, Finland, for computational resources and Esa-Pekka Kumpula for his help in measuring TEM images. The facilities and expertise of the DDCB Unit at the Faculty of Pharmacy, University of Helsinki, supported by HiLIFE and Biocenter Finland, is gratefully acknowledged.

## Author CRediT statement

Conceptualization: Artturi Koivuniemi

Data curation: Sirine Nouri, Akseli Niemelä

Formal analysis: Sirine Nouri

Funding acquisition: Artturi Koivuniemi

Investigation: Sirine Nouri, Akseli Niemelä, Sagar Dhakal, Laura Giorgi, Juho Heininen

Methodology: Sirine Nouri, Akseli Niemelä, Laura Giorgi, Khalfa Benadouda, Juho Heininen

Project administration: Sirine Nouri, Artturi Koivuniemi

Resources: Artturi Koivuniemi

Software: Sirine Nouri, Akseli Niemelä, Juho Heininen

Supervision: Sirine Nouri, Artturi Koivuniemi

Validation: Sirine Nouri

Visualization: Sirine Nouri

Writing – original draft: Sirine Nouri, Akseli Niemelä, Juho Heininen

Writing – review & editing: Sirine Nouri, Artturi Koivuniemi, Akseli Niemelä, Laura Giorgi, Juho Heininen

## References

1. Von Eckardstein A, Nofer JR, Assmann G. High Density Lipoproteins and Arteriosclerosis: Role of Cholesterol Efflux and Reverse Cholesterol Transport. Arterioscler Thromb Vasc Biol. 2001 Jan;21(1):13–27.

2. Pownall HJ, Rosales C, Gillard BK, Gotto AM. High-density lipoproteins, reverse cholesterol transport and atherogenesis. Nat Rev Cardiol. 2021 Oct;18(10):712–23.

3. Bhale AS, Meilhac O, d’Hellencourt CL, Vijayalakshmi MA, Venkataraman K. Cholesterol transport and beyond: Illuminating the versatile functions of HDL apolipoproteins through structural insights and functional implications. BioFactors. 2024 Sep;50(5):922–56.

4. Tall AR. Cholesterol efflux pathways and other potential mechanisms involved in the athero-protective effect of high density lipoproteins. J Intern Med. 2008 Mar;263(3):256–73.

5. Weber C, Noels H. Atherosclerosis: current pathogenesis and therapeutic options. Nat Med. 2011 Nov;17(11):1410–22.

6. Calabresi L, Franceschini G. Lecithin:Cholesterol Acyltransferase, High-Density Lipoproteins, and Atheroprotection in Humans. Trends Cardiovasc Med. 2010 Feb;20(2):50–3.

7. Zannis VI, Chroni A, Krieger M. Role of apoA-I, ABCA1, LCAT, and SR-BI in the biogenesis of HDL. J Mol Med. 2006 Apr;84(4):276–94.

8. Cao Y ni, Xu L, Han Y chun, Wang Y nan, Liu G, Qi R. Recombinant high-density lipoproteins and their use in cardiovascular diseases. Drug Discov Today. 2017 Jan;22(1):180–5.

9. Kornmueller, Vidakovic, Prassl. Artificial High Density Lipoprotein Nanoparticles in Cardiovascular Research. Molecules. 2019 Aug 2;24(15):2829.

10. Chen J, Zhang X, Millican R, Creutzmann JE, Martin S, Jun HW. High density lipoprotein mimicking nanoparticles for atherosclerosis. Nano Converg. 2020 Dec;7(1):6.

11. B. Uribe K, Benito-Vicente A, Martin C, Blanco-Vaca F, Rotllan N. (r)HDL in theranostics: how do we apply HDL’s biology for precision medicine in atherosclerosis management? Biomater Sci. 2021;9(9):3185–208.

12. Nicholls SJ, Andrews J, Kastelein JJP, Merkely B, Nissen SE, Ray KK, et al. Effect of Serial Infusions of CER-001, a Pre-β High-Density Lipoprotein Mimetic, on Coronary Atherosclerosis in Patients Following Acute Coronary Syndromes in the CER-001 Atherosclerosis Regression Acute Coronary Syndrome Trial: A Randomized Clinical Trial. JAMA Cardiol. 2018 Sep 1;3(9):815.

13. Gibson CM, Duffy D, Korjian S, Bahit MC, Chi G, Alexander JH, et al. Apolipoprotein A1 Infusions and Cardiovascular Outcomes after Acute Myocardial Infarction. N Engl J Med. 2024 May 2;390(17):1560–71.

14. Povsic TJ, Korjian S, Bahit MC, Chi G, Duffy D, Alexander JH, et al. Effect of Reconstituted Human Apolipoprotein A-I on Recurrent Ischemic Events in Survivors of Acute MI. J Am Coll Cardiol. 2024 Jun;83(22):2163–74.

15. Mathias RA, Velkoska E, Didichenko SA, Greene BH, Tan X, Navdaev AV, et al. Apolipoprotein A1 (CSL112) Increases Lecithin-Cholesterol Acyltransferase Levels in HDL Particles and Promotes Reverse Cholesterol Transport. JACC Basic Transl Sci. 2024 Nov;S2452302X24003978.

16. Navab M, Anantharamaiah GM, Reddy ST, Van Lenten BJ, Buga GM, Fogelman AM. Peptide mimetics of apolipoproteins improve HDL function. J Clin Lipidol. 2007 May;1(2):142–7.

17. Osei-Hwedieh DO, Amar M, Sviridov D, Remaley AT. Apolipoprotein mimetic peptides: Mechanisms of action as anti-atherogenic agents. Pharmacol Ther. 2011 Apr;130(1):83–91.

18. Sviridov D, Remaley AT. High-density lipoprotein mimetics: promises and challenges. Biochem J. 2015 Dec 15;472(3):249–59.

19. Anantharamaiah GM, Jones JL, Brouillette CG, Schmidt CF, Chung BH, Hughes TA, et al. Studies of synthetic peptide analogs of the amphipathic helix. Structure of complexes with dimyristoyl phosphatidylcholine. J Biol Chem. 1985 Aug;260(18):10248–55.

20. Datta G, Chaddha M, Hama S, Navab M, Fogelman AM, Garber DW, et al. Effects of increasing hydrophobicity on the physical-chemical and biological properties of a class A amphipathic helical peptide. J Lipid Res. 2001 Jul;42(7):1096–104.

21. Mishra VK, Palgunachari MN, Lund-Katz S, Phillips MC, Segrest JP, Anantharamaiah GM. Effect of the Arrangement of Tandem Repeating Units of Class A Amphipathic α-Helixes on Lipid Interaction. J Biol Chem. 1995 Jan;270(4):1602–11.

22. Wool GD, Reardon CA, Getz GS. Apolipoprotein A-I mimetic peptide helix number and helix linker influence potentially anti-atherogenic properties. J Lipid Res. 2008 Jun;49(6):1268– 83.

23. Sethi AA, Stonik JA, Thomas F, Demosky SJ, Amar M, Neufeld E, et al. Asymmetry in the Lipid Affinity of Bihelical Amphipathic Peptides. J Biol Chem. 2008 Nov;283(47):32273–82.

24. Dasseux JL, Sekul R, Bittner K, Cornut I, Metz G, Dufourcq J. Apolipoprotein A-I agonist and their use to treat dyslipidemic disorders. 6,004,925, 1999.

25. Esperion Therapeutics Inc. Esperion Reports Positive Results for Second Phase 1 Study of ETC-642. [Internet]. 2003. Available from: http://adisinsight.springer.com/downloads/mediarelease/1806/809029439.html

26. Anantharamaiah GM, Goldberg D, editors. Apolipoprotein Mimetics in the Management of Human Disease [Internet]. Cham: Springer International Publishing; 2015 [cited 2024 Mar 22]. Available from: https://link.springer.com/10.1007/978-3-319-17350-4

27. Benitez Amaro A, Solanelles Curco A, Garcia E, Julve J, Rives J, Benitez S, et al. Apolipoprotein and LRP1-Based Peptides as New Therapeutic Tools in Atherosclerosis. J Clin Med. 2021 Aug 13;10(16):3571.

28. Fawaz MV, Kim SY, Li D, Ming R, Xia Z, Olsen K, et al. Phospholipid Component Defines Pharmacokinetic and Pharmacodynamic Properties of Synthetic High-Density Lipoproteins. J Pharmacol Exp Ther. 2020 Feb;372(2):193–204.

29. Yuan W, Ernst K, Kuai R, Morin EE, Yu M, Sviridov DO, et al. Systematic evaluation of the effect of different apolipoprotein A-I mimetic peptides on the performance of synthetic high-density lipoproteins in vitro and in vivo. Nanomedicine Nanotechnol Biol Med. 2023 Feb;48:102646.

30. Wool GD, Vaisar T, Reardon CA, Getz GS. An apoA-I mimetic peptide containing a proline residue has greater in vivo HDL binding and anti-inflammatory ability than the 4F peptide. J Lipid Res. 2009 Sep;50(9):1889–900.

31. Zhao Y, Imura T, Leman LJ, Curtiss LK, Maryanoff BE, Ghadiri MR. Mimicry of High-Density Lipoprotein: Functional Peptide–Lipid Nanoparticles Based on Multivalent Peptide Constructs. J Am Chem Soc. 2013 Sep 11;135(36):13414–24.

32. Didichenko SA, Navdaev AV, Cukier AMO, Gille A, Schuetz P, Spycher MO, et al. Enhanced HDL Functionality in Small HDL Species Produced Upon Remodeling of HDL by Reconstituted HDL, CSL112: Effects on Cholesterol Efflux, Anti-Inflammatory and Antioxidative Activity. Circ Res. 2016 Sep 2;119(6):751–63.

33. Giorgi L, Niemelä A, Kumpula EP, Natri O, Parkkila P, Huiskonen JT, et al. Mechanistic Insights into the Activation of Lecithin–Cholesterol Acyltransferase in Therapeutic Nanodiscs Composed of Apolipoprotein A-I Mimetic Peptides and Phospholipids. Mol Pharm. 2022 Nov 7;19(11):4135–48.

34. Niemelä A, Koivuniemi A. Systematic evaluation of lecithin:cholesterol acyltransferase binding sites in apolipoproteins via peptide based nanodiscs: regulatory role of charged residues at positions 4 and 7. MacKerell A, editor. PLOS Comput Biol. 2024 May 28;20(5):e1012137.

35. Berendsen HJC, Van Der Spoel D, Van Drunen R. GROMACS: A message-passing parallel molecular dynamics implementation. Comput Phys Commun. 1995 Sep;91(1–3):43–56.

36. Maupetit J, Tuffery P, Derreumaux P. A coarse-grained protein force field for folding and structure prediction. Proteins Struct Funct Bioinforma. 2007 Nov;69(2):394–408.

37. Humphrey W, Dalke A, Schulten K. VMD: Visual molecular dynamics. J Mol Graph. 1996 Feb;14(1):33–8.

38. Sreerama N, Woody RW. Estimation of Protein Secondary Structure from Circular Dichroism Spectra: Comparison of CONTIN, SELCON, and CDSSTR Methods with an Expanded Reference Set. Anal Biochem. 2000 Dec;287(2):252–60.

39. Homan R, Esmaeil N, Mendelsohn L, Kato GJ. A fluorescence method to detect and quantitate sterol esterification by lecithin:cholesterol acyltransferase. Anal Biochem. 2013 Oct;441(1):80–6.

40. Gantt E. Properties and Ultrastructure of Phycoerythrin From *Porphyridium cruentum*. Plant Physiol. 1969 Nov 1;44(11):1629–38.

41. Pasti AP, Rossi V, Di Stefano G, Brigotti M, Hochkoeppler A. Human lactate dehydrogenase A undergoes allosteric transitions under pH conditions inducing the dissociation of the tetrameric enzyme. Biosci Rep. 2022 Jan 28;42(1):BSR20212654.

42. Nagazaki T, Akanuma Y. A new colorimteric method for the determination of plasma lecithin-cholesterol acyltransferase activity. Clin Chim Acta. 1977;75:371–5.

43. Calabresi L, Pisciotta L, Costantin A, Frigerio I, Eberini I, Alessandrini P, et al. The Molecular Basis of Lecithin:Cholesterol Acyltransferase Deficiency Syndromes: A Comprehensive Study of Molecular and Biochemical Findings in 13 Unrelated Italian Families. Arterioscler Thromb Vasc Biol. 2005 Sep;25(9):1972–8.

44. Gu F, Jones MK, Chen J, Patterson JC, Catte A, Jerome WG, et al. Structures of Discoidal High Density Lipoproteins. J Biol Chem. 2010 Feb;285(7):4652–65.

45. Manthei KA, Patra D, Wilson CJ, Fawaz MV, Piersimoni L, Shenkar JC, et al. Structural analysis of lecithin:cholesterol acyltransferase bound to high density lipoprotein particles. Commun Biol. 2020 Jan 15;3(1):28.

46. Gursky O. Structural stability and functional remodeling of high-density lipoproteins. FEBS Lett. 2015 Sep 14;589(19PartA):2627–39.

47. Al Musaimi O, Lombardi L, Williams DR, Albericio F. Strategies for Improving Peptide Stability and Delivery. Pharmaceuticals. 2022 Oct 19;15(10):1283.

48. Salnikov ES, Anantharamaiah GM, Bechinger B. Supramolecular Organization of Apolipoprotein-A-I-Derived Peptides within Disc-like Arrangements. Biophys J. 2018 Aug;115(3):467–77.

49. Islam RM, Pourmousa M, Sviridov D, Gordon SM, Neufeld EB, Freeman LA, et al. Structural properties of apolipoprotein A-I mimetic peptides that promote ABCA1-dependent cholesterol efflux. Sci Rep. 2018 Feb 13;8(1):2956.

50. Remaley AT, Thomas F, Stonik JA, Demosky SJ, Bark SE, Neufeld EB, et al. Synthetic amphipathic helical peptides promote lipid efflux from cells by an ABCA1-dependent and an ABCA1-independent pathway. J Lipid Res. 2003 Apr;44(4):828–36.

51. Natarajan P, Forte TM, Chu B, Phillips MC, Oram JF, Bielicki JK. Identification of an Apolipoprotein A-I Structural Element That Mediates Cellular Cholesterol Efflux and Stabilizes ATP Binding Cassette Transporter A1. J Biol Chem. 2004 Jun;279(23):24044–52.

52. Cavaco M, Andreu D, Castanho MARB. The Challenge of Peptide Proteolytic Stability Studies: Scarce Data, Difficult Readability, and the Need for Harmonization. Angew Chem. 2021 Jan 25;133(4):1710–2.

53. Graversen JH, Laurberg JM, Andersen MH, Falk E, Nieland J, Christensen J, et al. Trimerization of Apolipoprotein A-I Retards Plasma Clearance and Preserves Antiatherosclerotic Properties. J Cardiovasc Pharmacol. 2008 Feb;51(2):170–7.

54. Ditiatkovski M, Palsson J, Chin-Dusting J, Remaley AT, Sviridov D. Apolipoprotein A-I Mimetic Peptides: Discordance Between In Vitro and In Vivo Properties. 2017;

