## Supplemental information for "Modifications of the 22A apoA-I mimetic peptide sequence improve the anti-atherosclerotic properties of synthetic HDL"

### Detailed methods

#### *Molecular dynamics simulation*

Coarse-grained molecular dynamics simulations using the MARTINI 3.0 force-field (1) were run with the GROMACS software package (version 2020.2) (2). Parameters for DMPC and water were included in the 3.0 release of MARTINI. Initial coordinates for peptides were attained with PEPFOLD4 software (3) and for LCAT the protein preparation wizard of SCHRÖDINGER software package (Release 2020-3. Schrödinger, LLC, New York, 2021) was used to prepare PDB ID 6MVD (4). Standard settings in the protein preparation wizard were used, with missing loops fixed with Prime, protonation states generated and optimized with Epik and PROPKA using pH 7.0 and the final structure energy minimized with OPLS3e. The initial coordinates were fed to Martinize2 (version 0.9.6) to attain MARTINI parameters for peptides and LCAT (5). MKDSSP was used to define their secondary structures and the tertiary structure of LCAT was fixed with an elastic network using default settings (6).

Production systems utilized a v-rescale thermostat set to 310 K with a 1 ps time constant and an isotropic Parrinello-Rahman barostat set to 1 bar with a 12 ps time constant and a $3*10^{-4}$ $bar^{-1}$ compressibility. Pair lists were handled with Verlet every 20 steps, Van der Waals interactions with a potential shift set to a cut-off of 1.1 nm and electrostatic interactions with a reaction field set to a cut-off of 1.1 nm. The relative dielectric constant was set to 15. Periodic boundary conditions were set to all sides and the time step was set to 20 fs. For compressing the random lipid-peptide complex together the same settings were used except the barostat was set to semi-isotropic with a compressibility of $1*10^{-4}$ $bar^{-1}$ in the x/y direction and $3*10^{-4}$ $bar^{-1}$ in the z direction. The systems were run for 100 ps.

To analyse the peptide distribution around the nanodisc the trajectory was transformed in the following way, utilizing MDAnalysis and scikit-spatial python packages (7). For each frame a plane was fit to the centers of mass of DMPC molecules and the center of mass of LCAT was projected on the plane. The vector from the plane centroid to the projected LCAT point was defined as the X-axis.

The plane normal was defined as the Z-axis, and the cross product of X- and Z-axes defined the Y-axis. Since for each frame the direction of the plane normal is arbitrary, if the angle between the new normal and the normal of the previous frame was over 90 degrees the normal was inverted. As the nanodisc could not rotate over 90 degrees between frames (1 ns), this ensures the Z-axis faces the same direction in every frame. The centers of mass of the 22A fragments of each of the peptides were used to generate the peptide distribution heat maps. The bins were normalized by peptide and frame number within systems, and the colorbar by the maximum value between systems, enabling a direct comparison. Using the transformed trajectory, we analysed the peptides tendency to rotate decoupled from the overall nanodisc rotations with gmx rotacf included in the GROMACS package. Peptide backbone beads of residues 2 and 21 were used for this analysis.

Peptide occupancy as percentage of simulated time in the peptide binding site of LCAT was calculated with the following definition: the backbone bead of peptide residue 2 has to be within 1.2 nm of the backbone bead of LCAT residue 226; the backbone bead of peptide residue 11 has to be within 1.2 nm of the center of mass of the backbone beads of LCAT residues 48 and 237; and the backbone bead of peptide residue 20 has to be within 2.2 nm of the backbone bead of LCAT residue 48. Since the 18A fragment of 22A-P-18A also showed a tendency to bind to the site, peptide residues 24, 33 and 41 were used as well respectively.

Peptide contacts to LCAT and LCAT contacts to peptides were calculated using gmx pairdist included in the GROMACS package. For peptide contacts to LCAT the beads at the same position on different peptide molecules were grouped together and all LCAT beads were grouped together, while for LCAT contacts to peptides, all peptide beads were grouped together. A contact was registered when the minimum distance between the groups was less that 0.6 nm.

*Stability of peptide in human plasma*

The LC-HRMS measurements were carried out using Waters Xevo QTOF mass spectrometer coupled to a Waters Acquity UPLC system. Chromatography was performed using Waters Acquity UPLC $C_{18}$ column (BEH C18 1.7 µm, 100 mm × 2.1 mm) and the column was held at held at 40 °C. The autosampler temperature was 8°C and the injection volume was 2 µL using partial loop with needle overfill injection mode. The mobile phase consisted of (A) water and (B) methanol (LC-MS chromasolv, Honeywell), both containing 0.1 % formic acid. The LC gradient started at 5% B and this was increased to 50% B within 3 minutes, then further to 90% B within the next 3 minutes. The gradient was held at 90% for 2.5 minutes and lowered to 5% B within next 0.5 minutes. After that, the gradient was equilibrated at 5% for 1 minute resulting in total gradient time of 10 min. The gradient was run using 0.3 mL/min flow rate.

The mass spectra were measured using electrospray ionization in positive mode with capillary voltage of 1.5 kV. The mass range was m/z 120-1500, scan time 0.3 second and internal mass calibration was performed using leucine enkephalin. The source temperature was set to 135 °C, desolvation temperature was set to 450 °C, and desolvation gas flow rate was set to 900 L/h. LC-HRMS optimization and analyte specificity were evaluated using analyte-free matrix and 50 µg/mL standard stock solutions. Analyte ions were measured with below 30 ppm mass accuracy. The most abundant ion among the observed charge states was used for quantification (Table S3) by integrating the extracted ion chromatogram with a 0.08 Da window using the Integrate-function in MassLynx. Additional analyte charge state ions were used as qualifiers.

**References**


1. Souza PCT, Alessandri R, Barnoud J, Thallmair S, Faustino I, Grünewald F, et al. Martini 3: a general purpose force field for coarse-grained molecular dynamics. Nat Methods. 2021 Apr;18(4):382–8.

2. Berendsen HJC, Van Der Spoel D, Van Drunen R. GROMACS: A message-passing parallel molecular dynamics implementation. Computer Physics Communications. 1995 Sep;91(1–3):43–56.

3. Rey J, Murail S, de Vries S, Derreumaux P, Tuffery P. PEP-FOLD4: a pH-dependent force field for peptide structure prediction in aqueous solution. Nucleic Acids Research. 2023 Jul 5;51(W1):W432–7.

4. Manthei KA, Yang SM, Baljinnyam B, Chang L, Glukhova A, Yuan W, et al. Molecular basis for activation of lecithin:cholesterol acyltransferase by a compound that increases HDL cholesterol. eLife. 2018 Nov 27;7:e41604.

5. Kroon P, Grunewald F, Barnoud J, Van Tilburg M, Souza P, Wassenaar T, et al. Martinize2 and Vermouth: Unified Framework for Topology Generation [Internet]. 2024 [cited 2025 Jun 4]. Available from: https://elifesciences.org/reviewed-preprints/90627v2

6. Touw WG, Baakman C, Black J, te Beek TAH, Krieger E, Joosten RP, et al. A series of PDB-related databanks for everyday needs. Nucleic Acids Research. 2015 Jan 28;43(D1):D364–8.

7. Michaud-Agrawal N, Denning EJ, Woolf TB, Beckstein O. MDAnalysis: A toolkit for the analysis of molecular dynamics simulations. J Comput Chem. 2011 Jul 30;32(10):2319–27.

| System | Top 10 peptide beads in contact with any LCAT bead | Fraction of simulation time < 0.6 nm to any LCAT bead | Top 10 LCAT beads in contact with any peptide bead | Fraction of simulation time < 0.6 nm to any peptide bead |
|---|---|---|---|---|
| 22A | 4-ASP-SC1 | 0.96 ± 0.03 | 70-LEU-BB | 0.71 ± 0.03 |
| | 7-ARG-SC2 | 0.96 ± 0.02 | 70-LEU-SC1 | 0.70 ± 0.03 |
| | 1-PRO-BB | 0.93 ± 0.03 | 73-ASP-SC1 | 0.69 ± 0.03 |
| | 3-LEU-SC1 | 0.91 ± 0.04 | 48-TRP-SC3 | 0.69 ± 0.01 |
| | 3-LEU-BB | 0.91 ± 0.05 | 72-VAL-SC1 | 0.68 ± 0.04 |
| | 7-ARG-SC1 | 0.89 ± 0.04 | 231-ILE-BB | 0.67 ± 0.05 |
| | 4-ASP-BB | 0.89 ± 0.04 | 236-SER-SC1 | 0.67 ± 0.03 |
| | 8-GLU-SC1 | 0.88 ± 0.02 | 48-TRP-SC4 | 0.67 ± 0.02 |
| | 1-PRO-SC1 | 0.84 ± 0.04 | 227-ASP-BB | 0.66 ± 0.05 |
| | 2-VAL-BB | 0.84 ± 0.06 | 48-TRP-SC5 | 0.66 ± 0.02 |
| 22A-F | 4-ASP-SC1 | 0.93 ± 0.02 | 231-ILE-BB | 0.69 ± 0.04 |
| | 7-ARG-SC2 | 0.91 ± 0.01 | 227-ASP-BB | 0.69 ± 0.03 |
| | 1-PRO-BB | 0.89 ± 0.03 | 335-ASP-SC1 | 0.68 ± 0.02 |
| | 3-LEU-SC1 | 0.89 ± 0.00 | 48-TRP-SC3 | 0.66 ± 0.01 |
| | 3-LEU-BB | 0.89 ± 0.01 | 236-SER-SC1 | 0.66 ± 0.03 |
| | 22-LYS-SC2 | 0.88 ± 0.02 | 227-ASP-SC1 | 0.65 ± 0.00 |
| | 4-ASP-BB | 0.83 ± 0.03 | 70-LEU-SC1 | 0.65 ± 0.02 |
| | 7-ARG-SC1 | 0.82 ± 0.01 | 72-VAL-SC1 | 0.64 ± 0.05 |
| | 2-VAL-BB | 0.82 ± 0.02 | 379-ASN-SC1 | 0.64 ± 0.04 |
| | 1-PRO-SC1 | 0.81 ± 0.04 | 48-TRP-SC4 | 0.64 ± 0.01 |
| 22A-P-18A | 24-ASP-SC1 | 0.90 ± 0.05 | 335-ASP-SC1 | 0.78 ± 0.04 |
| | 22-LYS-SC2 | 0.89 ± 0.04 | 231-ILE-BB | 0.73 ± 0.02 |
| | 22-LYS-SC1 | 0.83 ± 0.06 | 227-ASP-BB | 0.72 ± 0.01 |
| | 24-ASP-BB | 0.81 ± 0.05 | 229-GLN-BB | 0.72 ± 0.03 |
| | 20-LYS-SC2 | 0.79 ± 0.04 | 48-TRP-SC3 | 0.70 ± 0.02 |
| | 25-TRP-SC3 | 0.77 ± 0.03 | 227-ASP-SC1 | 0.69 ± 0.04 |
| | 25-TRP-SC2 | 0.77 ± 0.03 | 70-LEU-BB | 0.69 ± 0.05 |
| | 27-LYS-SC2 | 0.77 ± 0.09 | 70-LEU-SC1 | 0.68 ± 0.04 |
| | 28-ALA-SC1 | 0.76 ± 0.03 | 48-TRP-SC4 | 0.68 ± 0.02 |
| | 25-TRP-SC5 | 0.76 ± 0.03 | 379-ASN-SC1 | 0.67 ± 0.02 |

Table S1: Contact analysis of molecular dynamics simulations. The beads are named in the following way: “residue number-amino acid identifier-bead type”, where bead type is either backbone (BB) or side chain (SC). Only the bead with the 10 highest contacts are shown. When peptide contacts to LCAT were analyzed the peptide beads at the same position of different molecules were grouped together. In other words, a peptide bead was not in contact with LCAT only if that bead on all peptides was farther than 0.6 nm from all LCAT beads. Values are presented as mean ± SD of triplicate simulations.

| | | Dunnett's test p-value compared to 22A-sHDL | | |
|---|---|---|---|---|
| | | 22A-F-sHDL | 22A-P-18A-sHDL | DMPC control |
| Cholesterol efflux values at different sHDL concentrations (μM) | 3 | 0.262 | 0.003 | 0.884 |
| | 6 | 0.093 | 0.001 | 0,408 |
| | 10 | 0.022 | 0.003 | 0,985 |
| | 15 | 0.055 | 0.017 | 0,041 |
| | 20 | < 0.001 | 0.001 | < 0.001 |
| $K_{cat}$ values | | 0.881 | 0.008 | N/A |

Table S2: Statistical analysis of cholesterol efflux assay results and the enzymatic reaction between recombinant LCAT and DHE-containing sHDL particles. The cholesterol efflux percentages at each concentration point and the $K_{cat}$ values were compared to 22A-sHDL by one-way ANOVA with a post-hoc Dunnett's test. For cholesterol efflux values, the degrees of freedom of the comparisons were 3 and 8 for between groups and within groups respectively. For $K_{cat}$ values, the degrees of freedom of the comparisons were 2 and 6 for between groups and within groups respectively.

| Peptide | Ion | Exact mass | Role | Retention time (min) | Measured mass | Mass error (ppm) |
|---|---|---|---|---|---|---|
| 22A | $[M+2H]^{2+}$ | 1311.7838 | qualifier | 6.14 | 1311.8060 | 16.7 |
| | $[M+3H]^{3+}$ | 874.8585 | qualifier | 6.14 | 874.8644 | 6.7 |
| | $[M+4H]^{4+}$ | 656.3958 | qualifier | 6.14 | 656.3887 | 10.8 |
| | **$[M+5H]^{5+}$** | **525.3182** | **quantifier** | 6.14 | 525.3027 | 29.5 |
| | $[M+6H]^{6+}$ | 437.9332 | qualifier | 6.14 | 437.9224 | 24.7 |
| 22A-F | $[M+2H]^{2+}$ | 1385.3180 | qualifier | 6.24 | 1385.3540 | 26.3 |
| | $[M+3H]^{3+}$ | 923.8813 | qualifier | 6.24 | 923.8882 | 7.5 |
| | $[M+4H]^{4+}$ | 693.1629 | qualifier | 6.24 | 693.1545 | 12.1 |
| | **$[M+5H]^{5+}$** | **554.7319** | **quantifier** | 6.24 | 554.7169 | 27.0 |
| | $[M+6H]^{6+}$ | 462.4446 | qualifier | 6.24 | 462.4332 | 24.7 |
| 22A-P-18A | $[M+4H]^{4+}$ | 1226.1956 | qualifier | 6.06 | 1226.1970 | 1.1 |
| | $[M+5H]^{5+}$ | 981.1580 | qualifier | 6.06 | 981.1627 | 4.8 |
| | $[M+6H]^{6+}$ | 817.7997 | qualifier | 6.06 | 817.8068 | 8.7 |
| | **$[M+7H]^{7+}$** | **701.1151** | **quantifier** | 6.06 | 701.1127 | 3.4 |
| | $[M+8H]^{8+}$ | 613.6017 | qualifier | 6.06 | 613.5949 | 11.1 |
| | $[M+9H]^{9+}$ | 545.5357 | qualifier | 6.06 | 545.5320 | 6.8 |

Table S3. Measured peptide ions by LC-HRMS, including their retention times, exact and measured (accurate) masses, mass errors, and designation as quantifier or qualifier ions. Ions used for quantification are shown in bold.

| | Dunnett's test p-value compared to control |
|---|---|
| 22A | 0.981 |
| 22A-sHDL | 0.054 |
| 22A-F | 1.000 |
| 22A-F-sHDL | 0.053 |
| 22A-P-18A | 0.973 |
| 22A-P-18A-sHDL | 0.002 |

Table S4: Statistical analysis of LCAT activity results in human plasma. The cholesterol esterification rate values were compared to the control (basal activity of endogenous LCAT in human plasma) by one-way ANOVA with a post-hoc Dunnett's test. The degrees of freedom of the comparisons were 6 and 14 for between groups and within groups respectively.

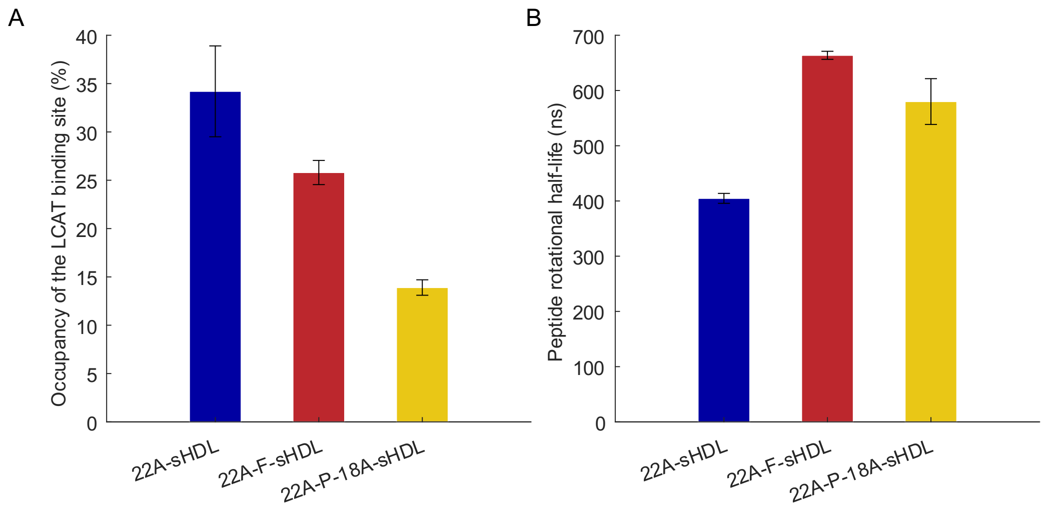


Figure S1: Molecular dynamics simulation analysis parameters. (A) Temporal occupancy percentages of peptides in the peptide binding site of LCAT. (B) Peptide rotational correlation function half-lives. Data are presented as mean ± SD.

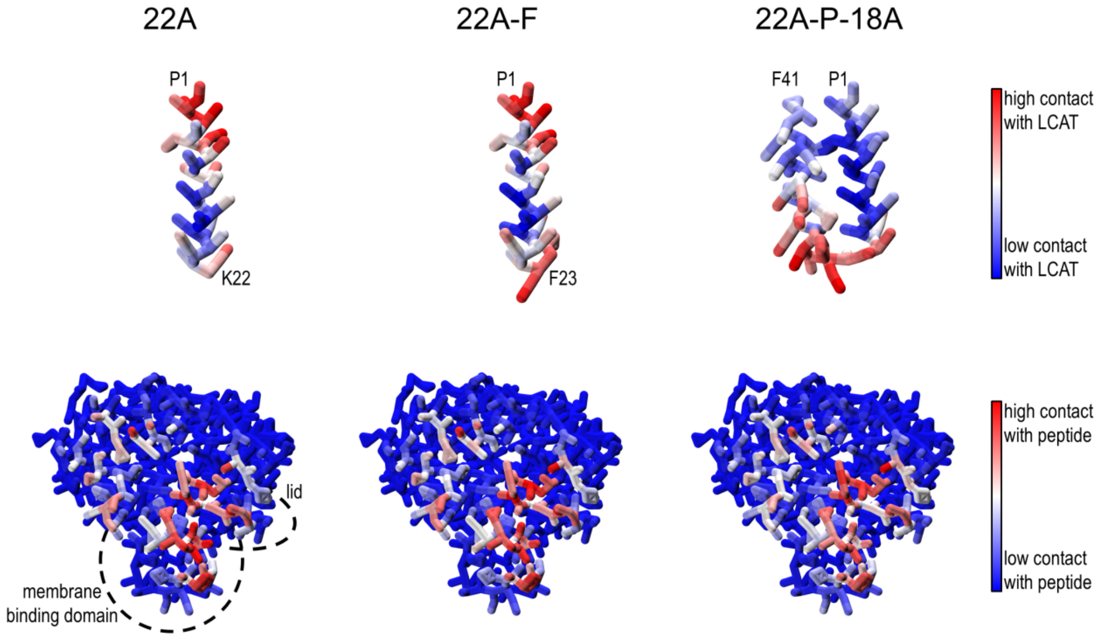


Figure S2: Contact analysis of molecular dynamics simulations visualized. The beads are colorized according to the mean value of triplicate simulations and the colorization normalized per system and per analysis.

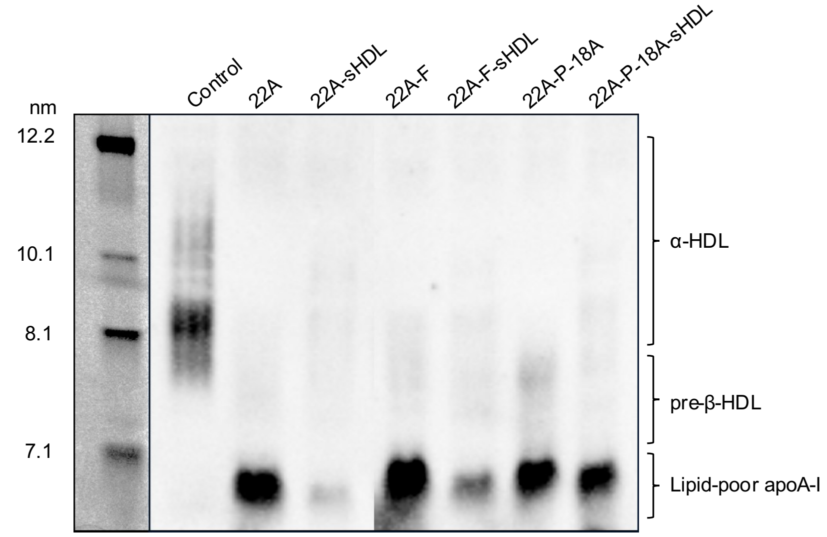


Figure S3: Remodeling of endogenous HDL induced by sHDL incubation with human plasma, duplicate assay. Plasma samples were incubated with different lipid-free peptides and sHDL particles for 1 hour at 37°C. Lipoproteins were separated by nondenaturing polyacrylamide gradient gel electrophoresis followed by Western blotting with an anti-apoA-I antibody. Two lanes were cut from the image (between 22A-sHDL and 22A-F) as these data corresponded to a peptide that is not included in this manuscript.
